# Longitudinal metatranscriptomic analysis of a meat spoilage microbiome detects abundant continued fermentation and environmental stress responses during shelf life and beyond

**DOI:** 10.1101/2020.08.13.250449

**Authors:** Jenni Hultman, Per Johansson, Johanna Björkroth

## Abstract

Microbial food spoilage is a complex phenomenon associated with the succession of the specific spoilage organisms (SSO) over the course of time. We performed a longitudinal metatranscriptomic study on a modified atmosphere packaged (MAP) beef product to increase understanding of the longitudinal behavior of a spoilage microbiome during shelf life and onward. Based on the annotation of the mRNA reads, we recognized three stages related to the active microbiome that were descriptive for the sensory quality of the beef: acceptable product (AP), early spoilage (ES) and late spoilage (LS). Both the 16S RNA taxonomic assignments from the total RNA and functional annotations of the active genes showed that these stages were significantly different from each other. However, the functional gene annotations showed more pronounced difference than the taxonomy assignments. Psychrotrophic lactic acid bacteria (LAB) formed the core of the SSO according to the transcribed reads. *Leuconostoc* species were the most abundant active LAB throughout the study period, whereas the activity of Streptococcaceae (mainly *Lactococcus*) increased after the product was spoiled. In the beginning of the experiment, the community managed environmental stress by cold-shock responses which were followed by the expression of the genes involved in managing oxidative stress. Glycolysis, pentose phosphate pathway and pyruvate metabolism were active throughout the study at a relatively stable level. However, the proportional activity of the enzymes in these pathways changed over time. For example, acetate kinase activity was characteristic for the AP stage whereas formate C-acetyltransferase transcription was associated with spoilage.

**Importance:** It is generally known which organisms are the typical SSO in foods, whereas the actively transcribed genes and pathways during microbial succession are poorly understood. This knowledge is important since better approaches to food quality evaluation and shelf life determination are needed. Thus, we conducted this study to find longitudinal markers that are connected to quality deterioration. These kind of RNA markers could be used to develop novel type of rapid quality analysis tools in the future. New tools are needed since even though SSO can be detected and their concentrations determined using the current microbiological methods, results from these analyses cannot predict how close timewise a spoilage community is from production of clear sensory defects. Main reason for this is that the species composition of a spoilage community does not change dramatically during late shelf life, whereas the ongoing metabolic activities lead to the development of notable sensory deterioration.

## Introduction

There has been an increasing market trend for case-ready meat, i.e. meat that is processed, packaged, and labeled at a central meat processing facility and delivered to the retail store ready to be put directly into the meat case. From the microbial ecology perspective, these packages are man-made niches in which microbial succession leading to spoilage occurs under the selective pressures of cold and the modified atmosphere (MA) applied (1). In comparison to atmosphere, increased CO_2_ levels (>20%) are used to limit the growth of aerobic food spoilage organisms, whereas high O_2_ concentrations (70% to 80%) are needed to keep myoglobin oxygenated to ensure the red appealing color of beef.

Even though case-ready meat products have great market value, we know relatively little about which pathways are active in a developing spoilage community over the course of time during shelf life. The use of CO_2_-containing atmospheres combined with refrigeration results in a dominance of psychrotrophic LAB, *Brochotrix thermosphacta* and, to some extent, *Enterobacteriaceae* in MAP meat (1, 2). During the last years, 16S rRNA gene amplicon-based approaches have increased the knowledge of community diversity in meat spoilage microbiomes (1, 3–9) or microbiota associated with food production environments (4, 7). However, no attention has yet been given to the activities related to the succession of a food spoilage community during shelf life.

We recently studied system level responses of three SSO during growth *in vitr*o (10), i.e. *Leuconostoc gelidum* subsp. *gasicomitatum* (subsp. *gasicomitatum*), *Lactococcus piscium*, and *Paucilactobacillus oligofermentans,* by comparing their time course transcriptome profiles obtained during growth. The study revealed how these LAB employed different strategies to cope with the consequences of interspecies competition. The fastest-growing bacterium, subsp. *gasicomitatum*, enhanced its nutrient-scavenging and growth capabilities in the presence of other LAB through upregulation of carbohydrate catabolic pathways, pyruvate fermentation enzymes, and ribosomal proteins, whereas the slower-growing *L. piscium* and *P. oligofermentans* downregulated these functions (10). This current study prompted from the observations we made about the different behavior of the three LAB while growing as communities *in vitro* (10). We wanted to investigate if we can recognize similar phenomena in the behavior of the SSO growing on meat over the course of time.

To create a comprehensive view of the activities of the spoilage community over the course of time, we analyzed the metatranscriptomes of natural beef spoilage communities developing at +6°C. We wanted to show which metabolic pathways and defense responses are transcribed during microbial succession and spoilage stages in a commercial meat product. Our aim was to study if we can distinguish different stages related to the sensory deterioration using longitudinal metatranscriptomic analysis.

## Results

We followed the development of a microbial community for 11 days using packages originating from the same production lot of a commercial MAP beef product manufactured by a large-scale operator. Production lot refers to a daily production and consist over 1 000 kg of meat. The experiment timespan covered the shelf-life and two days beyon. Altogether, a total of 65.8 Gb of sequence data (413.7 M reads) were obtained from the samples (Supplementary table S1). Good quality RNA suitable for metatranscriptomic analyses was obtained from all samples from day 2 onwards. Alongside the total RNA and rRNA depleted RNA sequencing, the approaches included conventional microbiology, sensory and pH analyses associated with the meat quality.

### Bacterial levels and sensory analysis results

On the first sampling day, bacterial levels were typical for a fresh product (LOG 4.6 and LOG 4.5 CFU/g LAB and total aerobic count, respectively) and at the end of the experiment the concentrations had increased to LOG 8.3 and 8.2 CFU/g for LAB and total aerobic bacteria, respectively (Fig. 1.). During the 11 days, the pH of the meat dropped slightly, from 5.9 to 5.5 (Supplementary table S2) and the microbial metabolism had also changed the packaging gas composition: O_2_ content decreased from 74% to 57% and CO_2_ increased from 20.5% to 37%, respectively. These changes are indicative for the activity of psychrotrophic heteroferementative spoilage LAB as anticipated.

**Figure 1.**
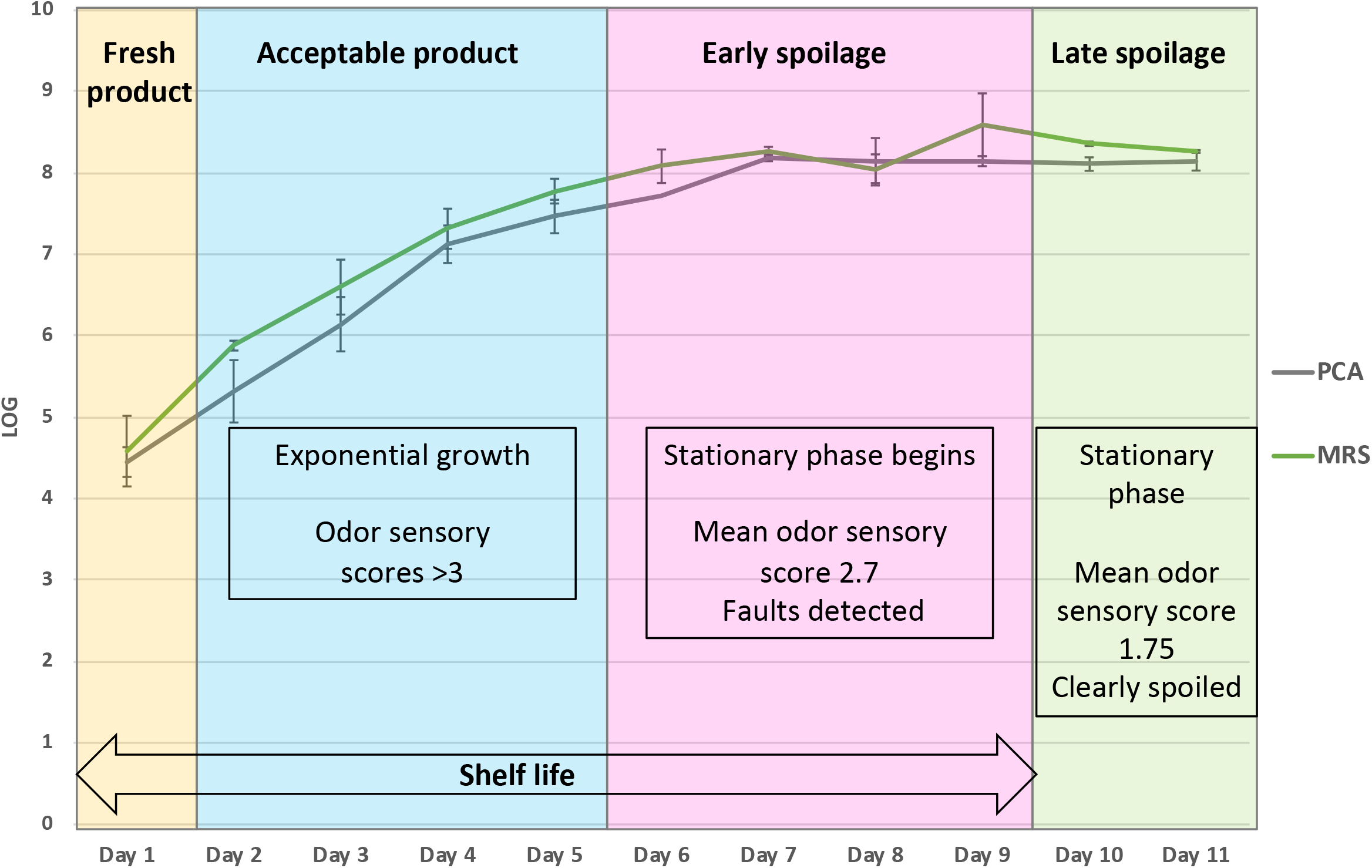
Bacterial concentrations (Plate count agar (PCA), total bacterial counts; de Mann Rogosa Sharpe agar (MRS), lactic acid bacterium counts) and main sensory findings during the experiment. The three stages related to sensory quality scores of the product were nominated as acceptable product (AP, days 3-5), early spoilage (ES, days 6-9), and late spoilage (LS, days 10-11) stages. Curve drawn from the mean values obtained from the results of two packages opened each day, whisker edges display concentration of each package.

For the bioinformatic analyses, the samples were divided into three groups based on the sensory analysis results (Fig. 1, Supplementary table S2) to enable detection of possible biomarkers associated with commercial meat quality during shelf life and beyond. The panelists started to deem the meat spoiled from day 5 on, with both odor and appearance receiving mean scores of 3/5. Samples assigned to the ES group (n=8 packages) received mean scores of 2.7/5 and 3.0/5 of odor and appearance, respectively. Samples receiving scores below 2 were assigned to the LS group and the mean odor and appearance scores were 1.75/5 and 1.65/5, respectively (Fig. 1, Supplementary table S2).

### Active microbial community composition during shelf life and beyond

Taxonomic assignment of the 16S rRNA gene fragments picked from the total RNA fraction indicated that the active microbial communities were composed mainly of *Firmicutes* from the families of Leuconostocaceae and Streptococcaceae, and unclassified bacilli (Fig. 2). During the first 5 days of the study (AP), Bacillales (unclassified bacilli and Bacillales) were active showing an abundance of 7.6% of the 16S rRNA gene transcripts but the abundance decreased beyond day 6 in ES. Leuconostocs were active throughout the experiment (Fig. 2) with abundance of 16.6-17.4%. In ES, the abundance of RNA transcripts of Streptococcaceae, mainly comprising sequences from the genus *Lactococcus*, increased to 5.2% of all detected 16S rRNA gene transcripts and this abundance remained high until the end of experiment, being 14.4% in LS. Based on the 16S rRNA results of the abundant genera and previous literature, we selected three SSO, i.e. subsp. *gasicomitatum* (LMG 18811^T^), *L. piscium* (MKFS47), *Dellaglioa algida* (DSM 15638^T^), for mapping of all reads against their genomes. The abundance of these psychrotrophic LAB was supported by the bowtie2 mapping since the majority of all RNA reads were assigned to these species (Table 1). The proportion of subsp*. gasicomitatum* was high from day 3 on (Table 1) whereas *L. piscium* reads increased on day 7 when the products had been deemed spoiled by the sensory panel. The reads mapping to *D. algida* showed highest abundance on day 4 (7.0 and 19.9%) but decreased during storage and thus this species was not very active during late shelf life and beyond (Table 1).

**Table 1.** Percentage of trimmed sequence reads from the mRNA fraction mapping to the genomes of three selected specific spoilage bacteria *Leuconostoc gelidum* subsp. *gasicomitatum* (LMG18811^T^), *Lactococcus piscium* (MKFS37) and *Dellaglioa algida* (DSM 15638^T^). Two packages were opened for each point of analysis.

| Sample | *L. gelidum* subsp.<br>*gasicomitatum* | *L. piscium* | *D. algida* | Sum |
| --- | --- | --- | --- | --- |
| Day2-1 | 3.2 | 0.1 | 1.1 | 4.4 |
| Day2-2 | 2.5 | 0.1 | 0.5 | 3.1 |
| Day3-1 | 19.7 | 0.7 | 7.1 | 27.5 |
| Day3-2 | 30.0 | 0.8 | 5.9 | 36.7 |
| Day4-1 | 27.7 | 3.9 | 7.0 | 38.6 |
| Day4-2 | 40.7 | 1.3 | 19.9 | 61.9 |
| Day5-1 | 43.7 | 1.7 | 6.4 | 51.8 |
| Day5-2 | 49.4 | 6.1 | 7.8 | 63,3 |
| Day6-1 | 39.7 | 5.9 | 4.8 | 50.4 |
| Day6-2 | 27.9 | 6.5 | 3.1 | 37.5 |
| Day7-1 | 20.5 | 17.4 | 4.2 | 42.1 |
| Day7-2 | 25.4 | 10.9 | 2.8 | 39.1 |
| Day8-1 | 37.8 | 10.4 | 5.4 | 53.6 |
| Day8-2 | 51.0 | 7.9 | 5.1 | 64.0 |
| Day9-1 | 15.5 | 8.7 | 4.0 | 28.2 |
| Day9-2 | 32.7 | 7.9 | 4.5 | 45.1 |
| Day10-1 | 13.2 | 2.4 | 1.7 | 17.3 |
| Day10-2 | 15.9 | 7.0 | 2.4 | 25.3 |
| Day11-1 | 18.6 | 4.0 | 1.4 | 24.0 |
| Day11-2 | 8.0 | 4.8 | 1.7 | 14.5 |

**Figure 2.**
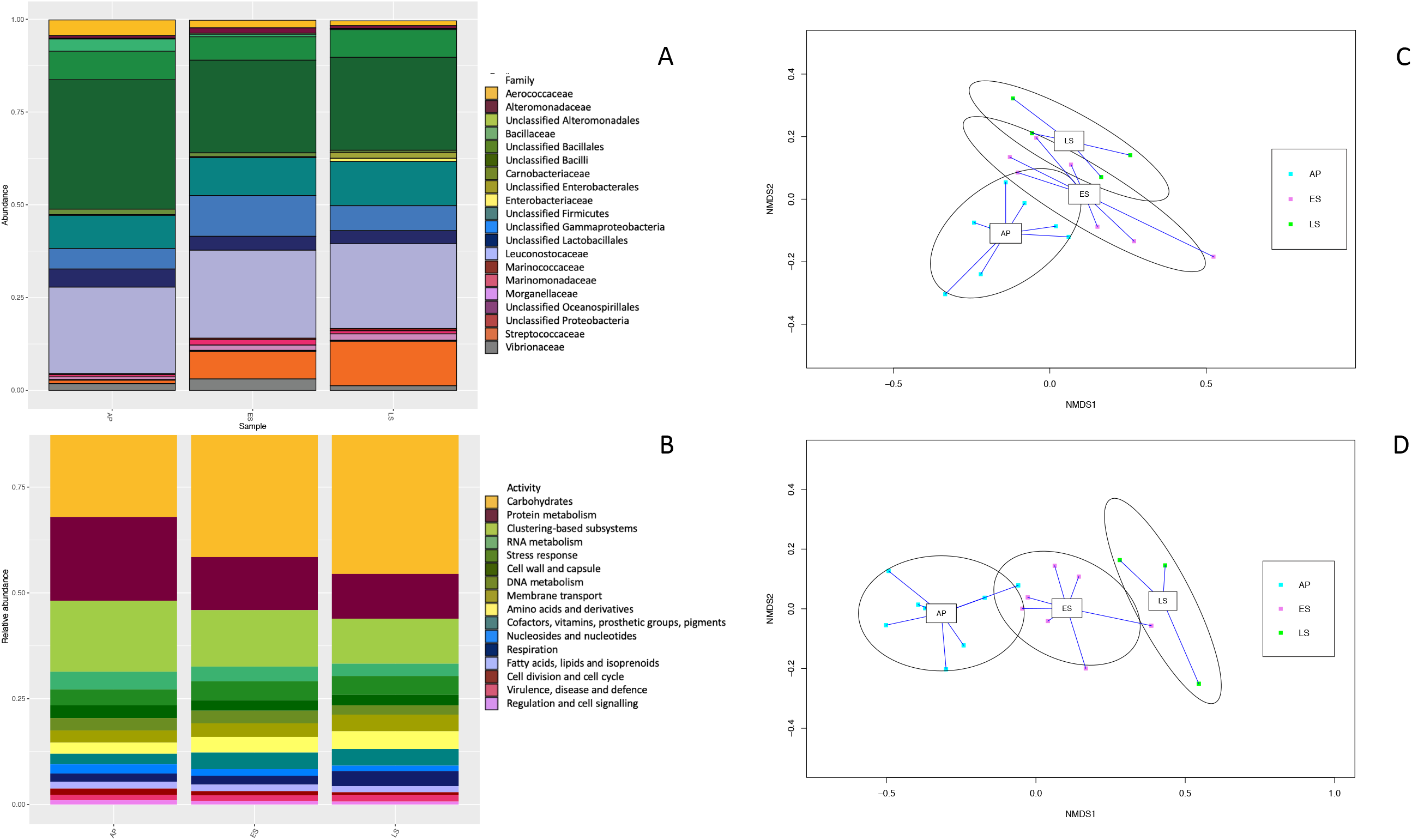
A) Assignment of 16S rRNA reads to taxa using Silva 138 database. The results of the most abundant families are shown. B) Change in activity during the experiment based on read annotation against SEED subsystems. C) Clustering of samples from different time points in nonmetric multidimensional scaling (NMDS) plot with 16S rRNA gene assignment and D) by the SEED database annotations. Samples from day 1 were discarded from the analysis due to the low number of reads recovered from sequencing. The samples were clustered into acceptable product (AP), early spoilage (ES) and late spoilage (LS). Blue lines show the labeled group centroids of the samples and the circle around the centroids is the 80% confidence area for standard deviation of the centroids.

### Three stages in the viewpoints of taxonomy and functional gene annotations

To explore the functional genes expressed over the course of time during shelf life, the sequenced mRNA fraction was analyzed. The results from read annotation against SEED subsystems database showed the community activity to change over time in the sample clusters related to stages AP, ES and LS. In NMDS plot of the Bray Curtis distance matrix, the samples clustered according to the stage. A permutation-based multivariate ANOVA (PERMANOVA) used for analyzing both the 16S rRNA gene taxonomic assignments and functional gene annotations revealed significant (R2=0.33, p<0.001) effect of the stage for taxonomic grouping of the samples (AP, ES, LS; Fig 2). However, the differences were more pronounced for the functional gene annotations (R2=0.67, p<0.01, Fig 2) than taxonomic assignments.

### Specific activities and changes in acceptable product

Microbial community at the acceptable product (AP) stage expressed cold shock genes (Fig. 3), indicating temperature stress at +6°C storage conditions. Genes related to protein biosynthesis constituted from 16% to 22% of all reads during exponential growth while decreasing to 12% and 9% in the later stages (Supplementary table S3). Active growth and succession of the community was thus strongly related to the stage while the product was still acceptable. According to SEED subsystems groups, cell division (1.1-1.25%, Supplementary table S3) was another function expressed significantly high during the exponential stage. These findings agree with the cultivation results indicating active community growth in the AP.

**Figure 3.**
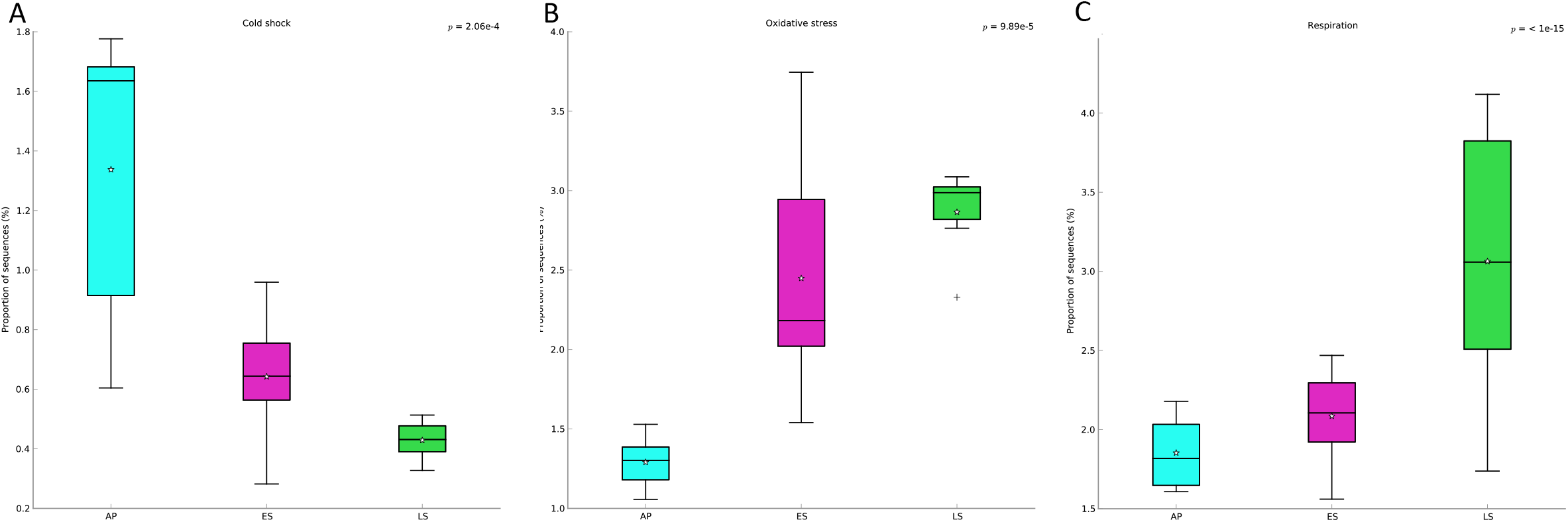
Relative abundance of transcripts involved in different SEED subsystem A) Cold shock B) Oxidative stress C) Respiration categories in AP, ES and LS with statistically differing abundances. Denote difference on y-axis in each of the plots. The median value is shown as line within the box and the mean value as a star.

During the AP, the abundance of gene transcripts clustering to “Carbohydrate metabolism” was one of the highest throughout the study and increased over time (Fig. 2). Detailed comparison of the carbohydrate metabolism in the three community growth stages showed that despite the rather similar expression level of the fermentation pathway, the relative abundance of genes transcribed in the pathways varied over time (Fig. 4, bubbles and pies related to glycolysis and pentose phosphate pathways). At the AP stage, the bacterial community required energy through fermentation via the pentose phosphate pathway (Fig. 4). Pyruvate metabolism was the second most abundant carbohydrate metabolism pathway during AP. However, unlike the spoilage stages, the majority of the transcripts were assigned as acetate kinase present in several different metabolic pathways. In the spoilage stages, the relative abundance of formate C-acetyltransferase, converting pyruvate to acetyl-CoA and formate, was significantly higher (Fig. 4.). Additionally, the third most abundant KO pathway was glycolysis, which had highly expressed genes throughout out the experiment (Fig. 4).

**Figure 4.**
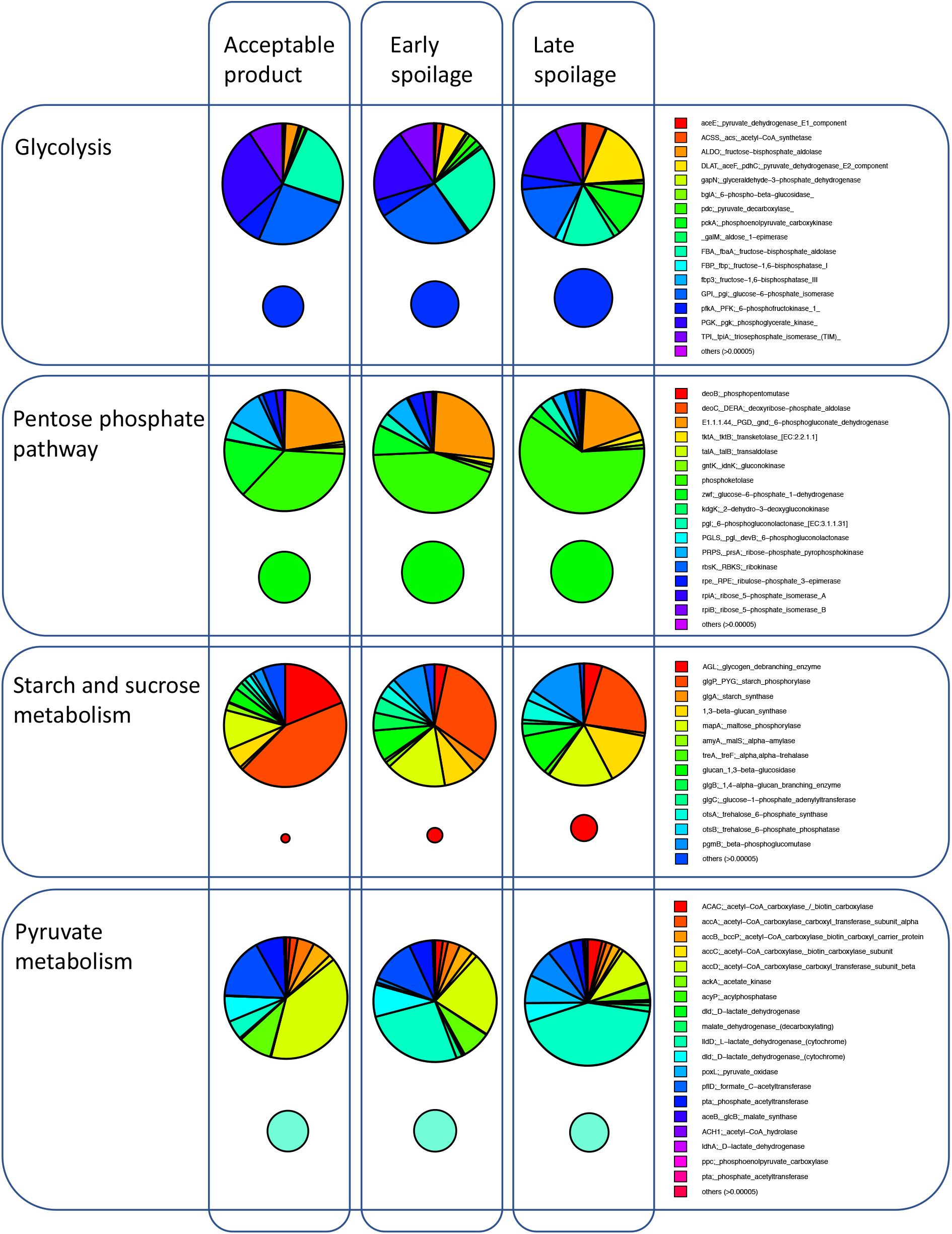
Carbohydrate metabolism activities of selected main pathways. Relative abundance of genes in pathways involved in carbohydrate metabolism in acceptable product (AP), early spoilage (ES) and late spoilage (LS) are denoted with the bubbles with sphere size relative to the abundance of transcripts. Pie charts visualize the abundance of expressed genes in each stage and the difference in gene transcript abundance. Most abundant genes of each pathway are shown and organized in clockwise order.

Mapping of the transcripts to the three selected SSO genomes showed that cold shock and universal stress pathways were active during the AP stage (Supplementary table S4). In subsp*. gasicomitatum,* the pentose phosphate pathway was expressed. In addition, amino acid synthesis was abundantly expressed. However, in *L. piscium,* genes related to lactate fermentation and glycolysis were expressed in relation to requirement of energy (Supplementary table S4).

### Specific activities and changes in samples showing early spoilage

After day 6 during the ES stage, the cold shock genes were no longer expressed (Fig. 3), and the succession had apparently led towards the cold-tolerant species and strains. The 16S rRNA data shows an increase in the proportion of psychrotrophic LAB with a decrease in mesophilic species. On the functional side there was a change as well: the abundance of transcripts involved in reaction to oxidative stress increased together with genes involved in microbial respiration (Fig. 3).

The members of the ES microbial community expressed genes involved in respiration as the activities of different cytochrome oxidases and ubiquinone menaquinone-cytochrome c reductase complexes (Supplementary table S3) acting in electron donating and accepting reactions were found to be abundant. The pathways with higher abundance in expression compared to the those in AP included formation of sugar alcohols, fermentation and central carbohydrate metabolism (Supplementary table S3). In more detail, the microbial communities expressed genes for glycolysis and gluconeogenesis and pentose phosphate pathway (Fig. 4, Supplementary table S3) as well as different fermentative energy requirement reactions (Supplementary table S3). Although stress response as a whole increased only slightly over time, the genes involved in oxidative stress were found more actively expressed from day 6 onward (Fig. 3). Similarly to cold shock reactions, the abundance of gene transcripts involved in cell division and cell cycle, motility and chemotaxis, nucleoside and nucleotide utilization and protein metabolism decreased with time (Supplementary table S3). This indicated that the community was no longer growing actively.

In the ES stage, subsp*. gasicomitatum* was abundant (48% based on *Leuconostoc* 16S rRNA gene abundance [Fig. 1] and 35% of the reads mapped to genome of subsp*. gasicomitatum* [Table 1]) and based on mapping of the RNA transcripts to the genome the active pathways included pentose phosphate pathway for energy requirement. The genes associated with stress and also the thioredoxin reductase activity increased compared to AP. Thioredoxin reductase has been shown to be related to survival in oxygen induced stress conditions (11) and this indicates that the species were actively using mechanisms to tolerate the stress caused by the high oxygen MAP. In addition, heme-dependent respiration of subsp*. gasicomitatum* (genes *cydA* and *cydB*) was found active in the AP and ES (Supplementary table S3).

At the ES stage, the second most abundant LAB species, *L. piscium,* utilized noxE NADH oxidases which can indicate a way to survive the redox stress in the high oxygen packaging during the active growth phase. Additionally, the aldehyde-alcohol dehydrogenases (adhE) reported to possess moonlight functions such as oxidative stress protection (12), were highly expressed (0.31+-0.21%). Unlike the heterofermentative subsp. *gasicomitatum,* which uses a phosphoketolase pathway for energy requirement, homoferementative *L. piscium* utilized glycolysis together with lactate and pyruvate fermentation (Supplementary table S4). Additionally, in ES the cold shock genes were highly active for *L. piscium* as well as the genes corresponding to sugar transporters. Moreover, in *L. piscium* gene *yngB*, coding for the fibronectin-binding protein was expressed at the ES stage (Supplementary table S4).

### Specific activities and changes associated with late spoilage

In pentose phosphate metabolism, the fermentation of xylulose to pentose (genes xfp, xpk; xylulose-5-phosphate/fructose-6-phosphate phosphoketolase) increased in the spoiled product, a phenomenon which is associated with the production of pyruvate (Supplementary table S3). In addition, there was increased activity of the pathway leading to the production of ethanol and acetate instead of common spoilage volatiles diacetyl and 2,3-butanediol. Similarly, the activity of the pentose metabolism gene *pflD* (formate C-acetyltransferase) increased in the LS stage. The gene catalyzes the reversible conversion of pyruvate into formate and is thus involved in the supply of pyruvate to the pentose phosphate pathway/metabolism.

The ES and LS stages differed in the utilization of di-and oligosaccharides and trehalose, which were both significantly higher in after use-by date samples. Trehalose acts as a stress protectant and storage carbohydrate and it is known that rapidly growing cells having lower trehalose levels than slow-growing and stationary-phase cells (13), explaining the lower activity in exponential phase cells. Malate metabolism increased significantly in the LS stage, especially the malate dehydrogenase, citrate synthase and isocitrate dehydrogenase genes associated with formation of citrate. During LS stage, starch metabolism was at its highest expression level (Fig. 4 bubbles) and also the relative abundance of genes transcribed in relation to it varied greatly over time (Fig. 4 pies).

In addition to carbohydrate metabolism, amino acid metabolism (arginine, urea cycle and polydeimines) increased with time as we found a seven-fold increase from ES to LS after the end-of shelf life stage (Supplementary table S3). These genes include *arcA*, *arcB*, *arcC*, *arcT* and *arcD*, corresponding to arginine deiminase, ornithine carbamouyltransferase, carbamate kinase, aspartate aminotransferase and arginine/ornithine antiporter.

After the product was spoiled at the LS stage, the majority of the reads mapping to subsp*. gasicomitatum* were involved in a pentose phosphate pathway producing pyruvate which, in the presence of oxygen, has been reported to lead to production of diacetyl (9, 24), a spoilage metabolite, through the chemical reaction of alpha acetolactate. As in ES, genes involved in general and oxygen-induced stress were active. However, unlike in ES, proteolytic genes (*clpL,* ATP-dependent Clp protease) became active and the abundance of reads mapping to sugar transporters increased. As was seen at the microbial community level, the trehalose metabolism increased at this stage and the genes were assigned to the main bacteria of this stage, subsp. *gasicomitatum*. For *L. piscium,* the metabolism did not change between ES and LS stages.

Based on the Metaphlann2 annotation (14) Ascomycota, especially yeasts from genus Debaryomycetaceae increased during shelf life (Fig. 5). The proportion of reads mapping to Debaryomycetaceae was 1.49% (±-2.37%) in the AP, 6.11% (±4.38%) in ES and increased to 25.62% (±12.74%) in LS. Based on the MG-RAST annotations, the reads from LS mapping to Debaryomycetaceae fungi had active energy metabolism/carbohydrate metabolism as genes involved in glycolysis, glyoxylate cycle and acetate metabolism were among those expressed (Supplementary table S5). Besides carbohydrate metabolism, fatty acid metabolism was active as the genes encoding glycerol-3-phosphate dehydrogenase, stearoyl-CoA desaturase and Acyl-CoA synthetase 2, to mention a few, had RNA transcripts mapping to them (Supplementary table S5). The activity of these yeasts was supported by finding RNA transcripts for several translation elongation factors as well as ribosomal 60S and 40S subunit ribosomal proteins. In yeasts, the genes involved in respiration were active too, and included different cytochrome C oxidases and oxioreductases as well as ferredoxin-dependent glutamate synthase. In addition, transaldolase, a gene from the non-oxidative phase of the pentose phosphate pathway, was active in yeasts during LS.

**Figure 5.**
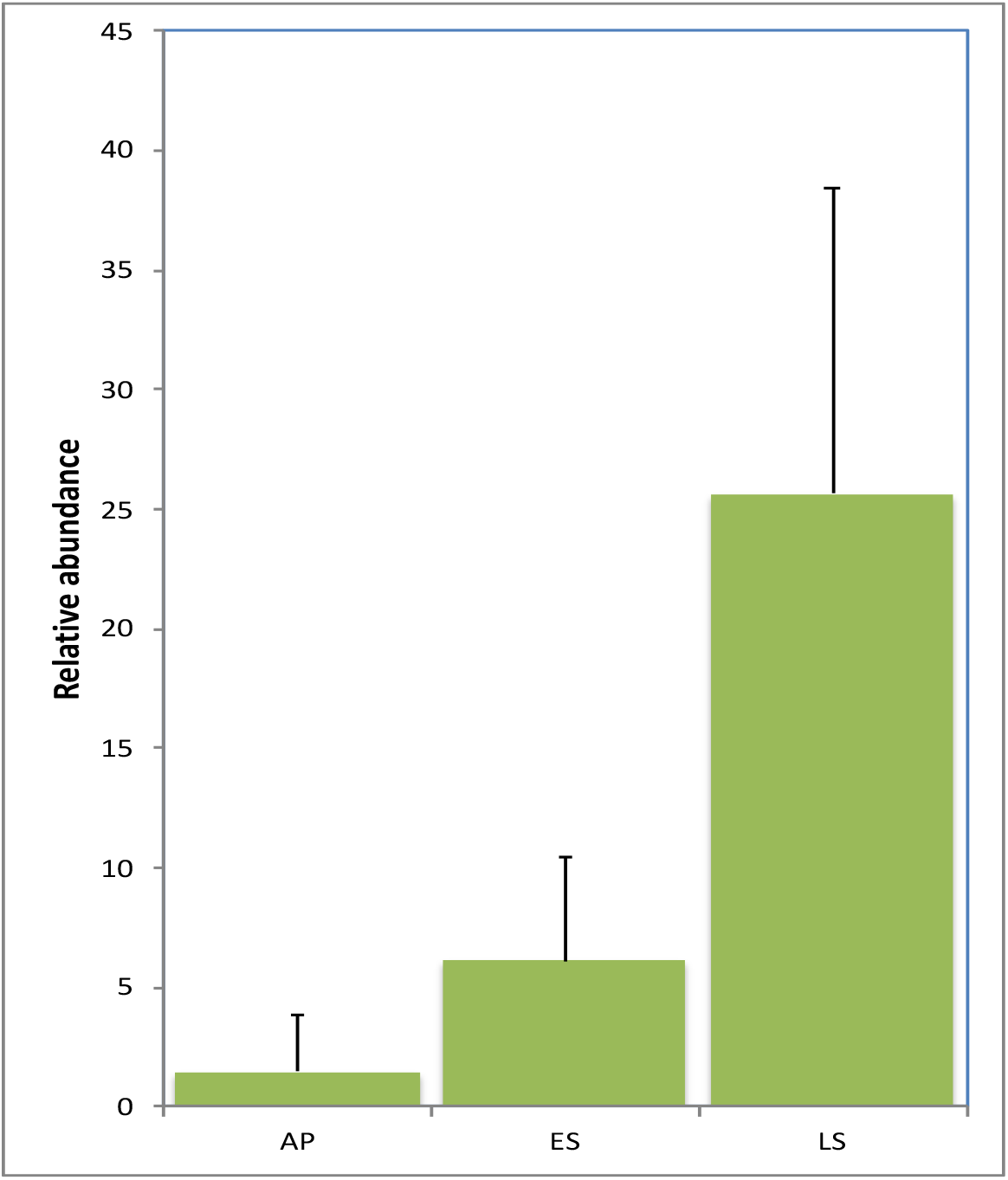
Proportion of reads belonging to ascomycotal family Debaryomycetaceae based on Humann2 annotation.

## Discussion

Based on the clustering of active transcripts in NMDS and the observed sensory characteristics and bacterial growth levels, the active microbiome and active genes in this MAP beef had three stages that were descriptive for the quality of the product: acceptable product (AP), early spoilage (ES) and late spoilage (LS). All these stages shared species and functions but there were several distinct metabolic and other responses associated with each of the three stages (Figs. 2 and 6). As typical for a packaged meat product (15), the microbial levels of the four stages did not directly correlate with the sensory quality of the product, whereas each of the four stages had specific activities based on the gene level activity analyses. Starting from the end of the AP stage, the same bacterial species remained active, something which highlights the reason why the active genes and pathways should be analyzed further instead of the active microbes to understand which metabolic pathways play a major role in food spoilage.

**Figure 6.**
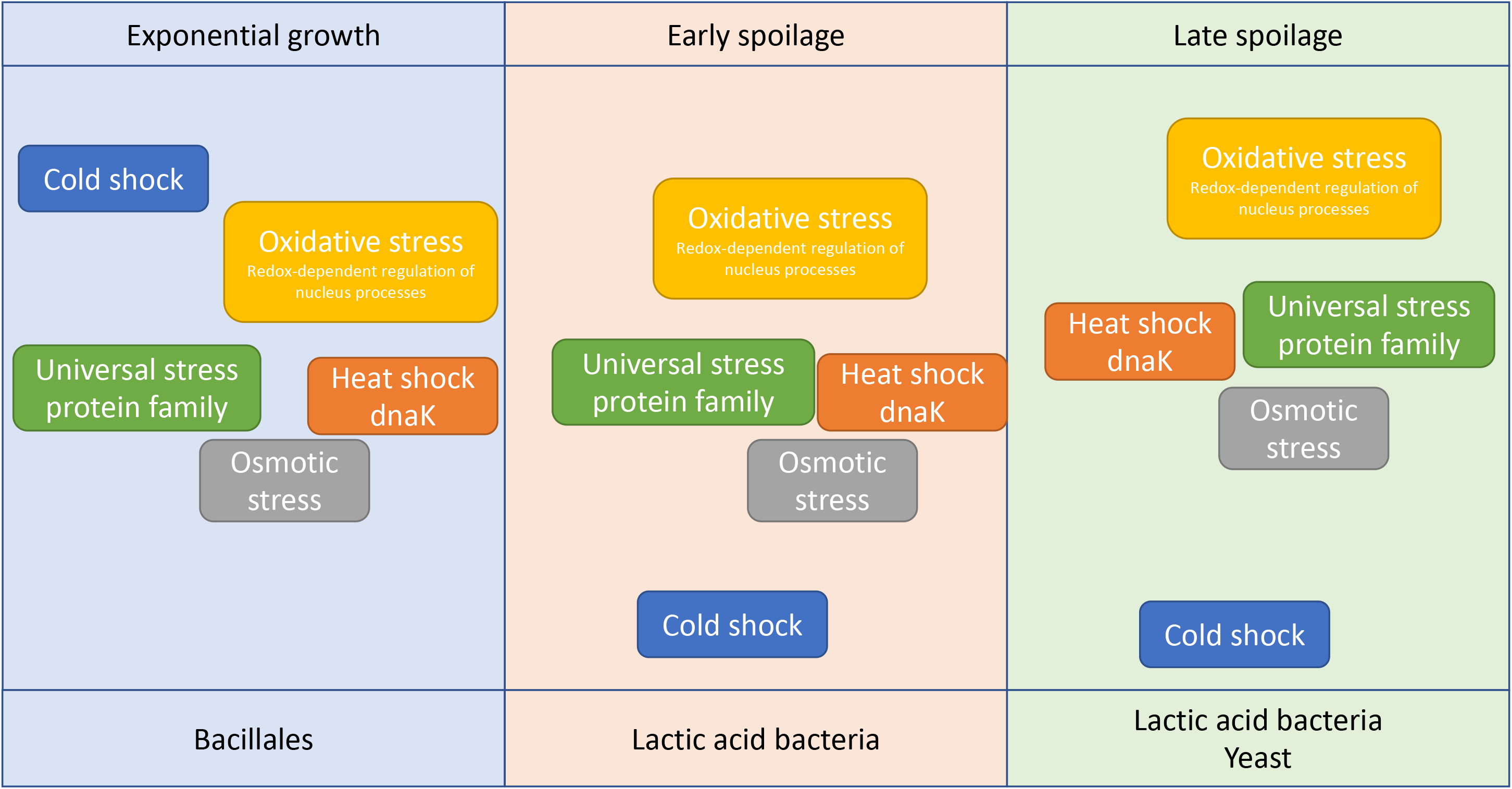
Most active functions and phyla in three stages, i.e. acceptable product, early spoilage and late spoilage, of a meat spoilage community developing in a high-oxygen modified atmosphere packaged beef product.

Even though leuconostocs, lactobacilli and lactococci have been associated with meat spoilage in several different studies using cultivation and DNA based approaches (16–19), little has been known about the most active functions during and after their succession. Based on the analysis of the active 16S rRNA genes in the total sequenced RNA fraction, leuconostocs were the most abundant bacteria throughout the study. However, the proportion or psychrotrophic *Lactococcus* increased after day 6. After shelf life (day 9), the LAB community was still active but the abundance of RNA transcripts of yeasts from the genus *Debaryomycetaceae* increased. Yeasts have not been considered to play a role in the spoilage of cold-stored meat before. Some cultivation-based studies have showed the fungi from *Saccharomycetaceae* phyla present in different meat environments, and also during the ripening of salami (20). The *Debaryomycetaceae* yeast increased drastically in the LS stage while the activity of the spoilage microbial community pertained to alcohol production from the pentose phosphate pathway. The yeasts were obviously active members of the fermentative community with up to 43.2% relative abundance in total microbial reads on the last day of the experiment (day 11) even though there was variation between different packages. Our results thus show that during long shelf-life, fungi and their spoilage functions increased in this MAP product.

Fermentation of carbohydrates was the most abundant metabolic activity detected in the samples over time as the prevalence of genes involved in carbohydrate metabolism was high throughout the study. The prevalence of actively expressed genes involved in carbohydrate metabolism shows that unlike in the spoilage of non-packaged meat caused by pseudomonads under atmosphere (21), the exhaustion of glucose is not a limiting factor for spoilage microbiota under MA containing 20% carbon dioxide and 80% oxygen. We observed activities of both homofermentative LAB, catabolizing glucose using the Embden–Meyerhof–Parnas (EMP) pathway, together with heterofermentative LAB, using the pentose phosphate or pentose phosphoketolase pathways. During the AP stage, when the community was in the exponential growth phase, pyruvate metabolism, especially the formate C-acetyltransferase, was expressed. In LAB capable of homofermentative metabolism, pyruvate formate lyase (PFL) is used during the transformation to mixed acid formation under glucose and oxygen limitation to increase the ATP yields (22). Here the beef was packaged under high oxygen MA and this enzyme is oxygen sensitive because it is cleaved and inhibited in the presence of oxygen (23). Nevertheless, the bacteria were found to express the *pfl* gene.

The high oxygen MA and storage at cold (+6° C) temperatures created stress for the bacterial communities (Fig. 3). The cold shock genes were found active during the first days of shelf life before the communities reached the exponential growth phase. However, in case of *L. piscium* cold shock genes were still expressed during ES but this species was growing delayed in comparison to subsp. *gasicomitatum*. In the lag phase, and especially after the commercially deemed shelf life was over, oxygen induced stress responses increased and thus the microbial community members had to strive to cope with the high oxygen MA. Heme-dependent respiration is among the ways to cope with oxygen related stress and also the relative abundance of transcripts involved in respiration increased. Based on previous functional genomics studies with single species (10, 18, 24), we hypothesized that in addition to the rapid onset of fermentation, the heme-dependent respiration of leuconostocs may play a major role for fitness under high oxygen MA also at the community level. Respiration has been shown to result in increased biomass of LAB, in several changes in the metabolism and in long-term survival (24–26). The increase in respiration with time indicates that oxygen stress becomes more evident for the community. Interestingly, respiration genes were found active in both bacterial and fungal members of the community in LS. Since the respiratory activity increased over time and was the highest in the AP and ES, the activity can be considered to play a role also in long-term survival and viability in a meat spoilage community during shelf life. In addition to the different cytochrome oxidases found in LAB, also ubiquinone Menaquinone-cytochrome c reductase complexes acting in electron donating and accepting reactions were active in the ES microbial community. In our recent coculture study (10), we noticed a similar trend in respect of subsp. *gasicomitatum*. It was the fastest-growing bacterium in the coculture containing *L. piscium* and *L. oligofermentans* (10). Subsp*. gasicomitatum* enhanced its nutrient-scavenging and growth capabilities in the presence of other LAB through upregulation of carbohydrate catabolic pathways, pyruvate fermentation enzymes, and ribosomal proteins (10). These findings are in line with the present study, highlighting the active role of this SSO in the community over the course of time. *D. algida*, a psychrotrophic LAB isolated from vacuum packaged beef, has been isolated from various beef products (27) but was not abundant in the beef product studied (Table 1). The 80% O_2_ clearly prevents growth of this bacteria and in a study where both high oxygen AMP and vacuum packaging were used, growth was higher in the latter (9). Thus, high oxygen MAP can be considered to support growth and activity of psychrotrophic LAB from the genera *Leuconostoc* and *Lactococcus*.

At the ES and LS stages, an increase in the activity of *noxE* gene was observed. As the microbes are growing actively while reaching the stationary phase, the use of carbohydrates leads to lack of potential electron acceptors. The *noxE* gene has been found to act in lowering the redox potential and thus enabling active metabolism (28). Based on the mapping, this gene was found active in *L. piscium* and thus this species can lower the redox potential of its environment to create more optimal growth conditions in MAP beef. The aldehyde-alcohol dehydrogenases were expressed at the ES and LS stages. In *E. coli,* this gene (*adhE*) has been shown to have antioxidant activity (29). Also, the *adhE* genes have been reported to possess moonlight functions such as oxidative stress protection (12, 29).

Gene *yngB* of *L. piscium,* encoding for fibronectin-binding protein, was actively transcribed in ES. This gene has been associated with heat stress in *Lactococcus lactis* but here we found indications of increased expression under oxidative stress on beef. In *L. piscium, yngB* (LACPI_1373, ([30)]) is a fibronectin and collagen binding protein, and it belongs to a class of fibronectin binding proteins without a signal peptide. Indications of the gene to be used both in fibronectin/collagen binding and biofilm formation have been documented for *S. suis* ortholog (31) and *Bacillus subtilis* (32).

In ES, the sugar alcohol metabolism increased. The substrate for metabolism may originate from the bacterial cell membranes since *Leuconostoc* and other LAB have genes for glycerophospholipid metabolism. Another possibility is that polyols produced by the yeast community increased the sugar alcohol metabolism (33).

In addition to carbohydrate metabolism and stress-induced reactions, we were able to capture novel information on the amino acid metabolism during meat spoilage. The arginine deimination is of interest since in *L. sakei* arginine deimination has been shown to increase the tolerance of both low pH and acid stress (34). Agmatine can be produced from arginine by a broad range of other bacteria through decarboxylation. The agamatine deiminase pathway, present e.g. in *L. piscium* (30), can help bacteria in protection against environmental stresses, including low pH in a similar way to how the arginine deiminase pathway does (35).

## Conclusions

Since sensory changes do not follow bacterial concentrations linearly, more accurate approaches for detecting food spoilage are needed. Results from our experiment show that instead of measuring bacterial cell numbers for product quality evaluation, the focus should be on more detailed activity markers related to food spoilage. These markers occurring over the course of time might provide a useful way to evaluate product freshness. A shift toward a *Lactococcus* dominated community can be considered as one indication of the end of shelf life for the meat product as well as the appearance of yeasts. In addition, the expression of genes involved in cold shock reactions decreased closer to the end of shelf life. Also, the activity of genes involved in respiration and oxygen stress were found to become more pronounced when the product was spoiled. Unlike what was previously thought, carbohydrate metabolism was one of the most active functions throughout the time and in the change from fresh to spoiled product.

## Materials and Methods

### Experiment description

Twenty-two packages of tenderized beef loin fillets stored in MAP were bought in commercial packaging. The samples were collected from the same lot (comprises production of one day) of a large-scale producer. During the experiment, the samples were stored at +6°C. Two packages were sampled destructively each day for 11 days. Before opening, the concentrations of O_2_ and CO_2_ in the package gas were measured (Checkpoint, PBI Dansensor) through a double septum and, after opening, one fillet (approximately 120 g) from each package was immediately transferred to a stomacher bag with 10 ml of RNA later (Ambion) and 5 ml of peptone water. The bacterial cells were removed from the beef surface by stomacher (Stomacher 400; Seward, West Sussex, United Kingdom) at a low power for 30 s. The liquid was collected from the stomacher bag and cell collection was conducted via two-stage centrifugation to separate the mammalian and bacterial cells. First, the supernatants were centrifuged in 15-ml conical tubes at 200 relative centrifugal force (RCF) for 3 min at +4°C to remove the fat and eukaryotic cells. The supernatant was collected and centrifuged in 1.5 ml Epperdorf tubes at 13000 rpm for 3 min at +4°C to collect the cells. The supernatant was poured off and replaced with 500 *μ*l of RNAlater. The collected cells were stored at −70°C until nucleic acid extraction.

Another fillet was used for quantitative microbiological and sensory analysis. Serial 10-fold dilution series were made from 22 g of beef with 198 ml of peptone-salt buffer (0.1% peptone, 0.9% NaCl) and homogenized by using a stomacher at medium power for 1 min, followed by cultivation onto MRS plates (LAB). In addition, plate count agar (PCA) was used to quantify the total cultivable microbes from the beef fillets. The PCA plates were incubated at 25°C for 3 to 5 days before colony enumeration. The MRS plates were incubated in jars made anaerobic by a commercial atmosphere generation system (AnaeroGen, Oxoid) at 25°C for 5 days, after which the CFU were calculated. Sensory analyses were performed by a trained panel of at least five individuals. For these analyses, the beef samples were equilibrated at room temperature. Beef samples from the same meat lot was stored fresh (day 0) in the freezer (− 20 °C) and used as a reference. The panelists evaluated the odor and the appearance of the samples using a five-point scale (1 = severe defect, spoiled; 2 = clear defect, spoiled; 3 = mild defect, satisfactory; 4 = good; 5 = excellent); the observed deficiencies were described by the panelists.

### RNA extraction and library preparation

The extraction protocol was modified from Chomczynski & Sacchi (36). First, the samples were centrifuged at 16 000 RCF at +4°C for 10 min and the RNAlater was carefully removed. The pellet was immediately covered with 500 *μ*l of Phenol-chloroform-isoamylalcohol (Sigma) and 500 *μ*l of denaturation solution (4M guanidium thicyanate, 24 mM sodium citrate, 0.5% sarcocyl and 0.1M betamercaptoethanol). The samples were transferred to Lysing matrix E tubes (mBio) and beated with beads in FastPrep-24 instrument (MP Biomedicals) for 30s at 5.5 m/s. After bead-beating, the tubes were placed on ice for 5 minutes followed by centrifugation at 13000 RCF at +4°C for 10 min. The supernatant was mixed with 500 ul chloroform, vortexed and centrifuged at 4°C at 13000 RCF for 5 min. Nucleic acids were precipitated with 1:10 of 3M NaAc, 10 × volume of isopropanol and 1 *μ*l of GlycoBlue (Invitrogen) at −20°C for 2 hours. The nucleic acid pellet was washed with 70% ethanol and eluted to 100 *μ*l of DEPC water. DNA and RNA were purified with Qiagen Allprep DNA RNA kit according to the manufacturer’s recommendations with additional DNAse treatment. Resulting DNA and RNA quality was checked with gel electrophoresis and Agilent Bioanalyzer Nano chips, respectively. Concentration was checked with Cubit (Thermo).

The sequencing libraries were made both from the ribosomal RNA (rRNA) depleted libraries (mRNA library) and total RNA. rRNA depletion for the mRNA library was conducted with Ribozero kit (Epicentre) with the low input protocol. Illumina TruSeq® Stranded mRNA kit was used the library preparation. The samples were sequenced on an Illumina NextSeq at the DNA sequencing and genomics laboratory at the University of Helsinki.

### Quality filtering

The raw reads were filtered for quality with cutadapt (37) to remove the reads with reverse adapter with quality cutoff Q20 and with a length below 60 bp. The quality filtered reads were mapped to *Bos taurus* genome (accession number NKLS00000000.2) using Bowtie2 (38) with default parameters and reads mapping to *B. taurus* were discarded.

### rRNA mapping

The rRNA reads from the quality filtered non-*B. taurus* reads of the total RNA libraries were searched with Metaxa2 (39) and blasted against Silva 138 database (40) to identify the 16S rRNA reads and thus analyze the change in active species composition of the samples.

### Annotation

All sequence reads from mRNA libraries were annotated against KEGG (41) and SEED (42) databases with e-value cutoff 1e-5, minimum identity cutoff 60% and min alignment cutoff 15 bp at MG-RAST (43). The annotated genes were normalized by sample read number. Three bacterial genomes *gelidum* subs. *gasicomitatum* (LMG 18811, FN822744.1), *L. piscium* (MKFS47, NZ_LN774769) and *D. algida* (DSM 15638, NZ_AZDI01000000) were selected for more thorough analysis. The sequence reads from all samples were mapped to the Uniprot (44) genes of each genome with Bowtie2 (38). In addition, the reads were annotated to species in Metaphlann2 (14) and specific pathways examined in Humann2 (45).

### Statistics

Bray-Curtis distance matrix from relative abundance normalized 16S rRNA picked from the total RNA libraries and metabolic pathway data from the mRNA libraries were ordinated with NMSD in R (46) using vegan (47), ggplot (48) and Phyloseq (49). Bacterial abundances were matched with the factor “Sample stage” with the function envfit from the Vegan package and 80% confidence area was used for standard deviation of the centroids. The statistical differences among the functional profiles of the three time points were analyzed with STAMP software package (50). An ANOVA analysis with Games-Howell post-hoc test and Storey’s FDR for correction was conducted. Sequence data were deposited at the European Nucleotide Archive (ENA) under the project PRJEB20288. The annotated metagenomes are available at MG-RAST under project BEEF_RNA.

## Acknowledgements

We thank Ms. Henna Niinivirta for skillful technical assistance. The CSC - IT Center for Science Ltd is acknowledged for providing computational resources. The project was financially supported by the Academy of Finland CODELAB (307855) and a grant to JH from the Walter Ehrström Foundation.

## Table 1 legend

**Table 1.** Percentage of trimmed sequence reads from the mRNA fraction mapping to the genomes of three selected specific spoilage bacteria *Leuconostoc gelidum* subsp. *gasicomitatum* (LMG18811^T^), *Lactococcus piscium* (MKFS37) and *Dellaglioa algida* (DSM 15638^T^). Two packages were opened for each point of analysis.

## Legends to the figures

**Figure 1**. Bacterial concentration (PCA: plate count agar, total bacterial counts; MRS: de Mann Rogosa Sharpe, LAB counts) and main sensory findings during experiment. The three stages related to the product were nominated as acceptable product (AP, days 3-5), early spoilage (ES, days 6-9), and late spoilage stage (LS, days 10-11). Curve presents mean value obtained from analyses of two packages, whiskers are related to each value obtained.

**Figure 2**. A) Assignment of 16S rRNA reads to taxa using Silva 138 database. The results of the most abundant families are shown. B) Change in activity during the experiment based on read annotation against SEED subsystems. C) Clustering of samples from different time points in nonmetric multidimensional scaling (NMDS) plot with 16S rRNA gene assignment and D) by the SEED database annotations. Samples from day 1 were discarded from the analysis due to the low number of reads recovered from sequencing. The samples were clustered into acceptable product (AP), early spoilage (ES) and late spoilage (LS). Blue lines show the labeled group centroids of the samples and the circle around the centroids is the 80% confidence area for standard deviation of the centroids.

**Figure 3**. Relative abundance of transcripts involved in different SEED subsystem A) Cold shock B) Oxidative stress C) Respiration categories in AP, ES and LS with statistically differing abundances. Denote difference on y-axis in each of the plots. The median value is shown as line within the box and the mean value as a star.

**Figure 4**. Carbohydrate metabolism activities of selected main pathways. Relative abundance of genes in pathways involved in carbohydrate metabolism in acceptable product (AP), early spoilage (ES) and late spoilage (LS) are denoted with the bubbles with sphere size relative to the abundance of transcripts. Pie charts visualize the abundance of expressed genes in each stage and the difference in gene transcript abundance. Most abundant genes of each pathway are shown and organized in clockwise order.

**Figure 5**. Proportion of reads belonging to ascomycotal family Debaryomycetaceae based on Humann2 annotation.

**Figure 6**. Most active functions and phyla in three stages, i.e. acceptable product, early spoilage and late spoilage, of a meat spoilage community developing in a high-oxygen modified atmosphere packaged beef product.

